# Biofilm formation by *Mycobacterium avium* ssp. *paratuberculosis* in aqueous extract of schmutzdecke for clarifying untreated water in water treatment operations

**DOI:** 10.1101/336370

**Authors:** G. Aboagye, M.T. Rowe

## Abstract

**AIMS:** To determine the effect of aqueous extract of schmutzdecke on adhesion and biofilm formation by three isolates of *Mycobacterium avium* ssp. *paratuberculosis* (Map) under laboratory conditions.

**METHODS AND RESULTS:** Strains of *Mycobacterium avium* ssp. *paratuberculosis* in aqueous extract of schmutzdecke were subjected to adhesion tests on two topologically different substrata i.e. aluminium and stainless steel coupons. Biofilm formation was then monitored in polyvinyl chloride (PVC) plates. All the three strains adhered onto both coupons, howbeit greatly on aluminium than stainless steel. In the PVC plates, however, all strains developed biofilms which were observed by spectrophotometric analysis.

**CONCLUSIONS:** The environmental isolates of Map attained higher cell proliferation in both filtered and unfiltered aqueous extracts of schmutzdecke (FAES and UAES respectively) compared with the human isolate. Furthermore, the results showed that irrespective of the media used, Map might have developed biofilm by its genetic competence to do so under favourable conditions that the immediate environment might have provided.

**Significance and impact of the study:** Composites of the schmutzdecke which is the dirty layer formed within 10 to 20 days of operation of a slow sand filter bed had a proliferative effect on Map. Therefore when entrapped, Map could form a biofilm and access human populations through potable water. Therefore, schmutzdecke should be monitored and scraped periodically to curtail its support for environmentally persistent pathogens that can pose health risks to humans.

**Highlights:**

- Schmutzdecke; a reservoir of nutrient composites atop slow sand filter bed
- Adhesion by Map on aluminium and stainless steel coupons was achieved
- Map strains from water sources developed biofilms better than the human strain
- In distilled water, biofilm formation by all strains was evident
- Protracted build-up of schmutzdecke may proliferate waterborne pathogens

## 1. INTRODUCTION

Mycobacteria possess a characteristic hydrophobic cell wall, a n d h a v e the genetic competence to form biofilms (Clary et al. 2018; Chern et al. 2015; Primm et al. 2004; Steed and Falkinham, 2006). They can attach and grow in biofilm on suitable substrata depending on growth requirements of the species involved, prevailing conditions in the environment during colonisation and the properties of the substrata (Hall-Stoodley et al. 2006). There have been many biofilm studies using a range of *Mycobacterium* species and inert and biological surfaces (Vaerewijck et al. 2005). Schulze-Röbbecke and Fischeder (1989) reported biofilm formation by *M. kansasii* and *M. flavescens* with cell densities of 2 x 10^5^ cfu cm^−2^ and 7 x 10^4^ cfu cm^−2^ respectively on the inner surface of silicone tubes after a period of 10 months. *Mycobacterium marinum* produced sinuous arrays of cells which formed into cords when grown on high-density polyethylene (HDPE), polycarbonate (PC) or silicon (Si) (Hall-Stoodley et al. 2006). Mycobacteria have also been demonstrated to form biofilm on biological substrata specifically by *M. ulcerans* forming biofilm in the salivary glands and legs of *Naucoris cimicoides* (Marsollier et al. 2005). Further evidences pertaining to mycobacterial biofilm formation have also been reported in aquatic environments. The non-tuberculous opportunistic pathogens of mycobacterial species have been recovered from biofilm associated with drinking water systems (September et al. 2004). Schulze-Röbbecke and Fischeder (1989) and Schulze-Röbbecke et al. (1992) recovered both slow and rapidly growing mycobacteria in 45 of 50 biofilm samples taken from municipal or domestic water supplies in Germany and France. The report by Falkinham et al. (2001) that treatment of water only had between 2 to 4 logs reduction effects on *M. avium* may have resulted from their persistence and biofilm formation capabilities (Primm et al. 2004). This evidence clearly demonstrates the possibility of mycobacterial species and the ability of *M. avium* and Map to form biofilm in water and agricultural environments (Lehtola et al. 2006; Beumer et al. 2010; Cook et al. 2010). Pickup et al. (2006) using a PCR assay based on the IS*900* insertion element found Map in biofilm on Nantgaredig Bridge spanning the River Tywi in Wales (UK) and on an abstraction site grating. There is therefore clear evidence, that mycobacteria are invariable residents of biofilm in water distribution systems. The presence of organic sediment, with which Map is presumably associated as part of the biofilm microflora, is an important feature in the persistence of the organism (Manning, 2001; Winterhoff et al. 2002) in lakes and rivers, and in the case of water treatment systems, the schmutzdecke which is a meshwork of biologically active matter atop the slow sand filtration may either promote or suppress the growth of the organism. Information regarding biofilm formation by Map is lacking but it does have a lipid cell wall (Wu et al. 2009) which contains on the cell envelope glycopeptidolipids (GPLs) which convey hydrophobic properties (Tatchou-Nyamsi-Konig et al. 2009). This makes Map potentially capable of attaching to a suitable surface, as GPLs are directly or indirectly required for colonisation of some surfaces (Freeman et al. 2006). Many different materials support biofilm formation by mycobacteria; high-density polyethylene (HDPE), polycarbonate and silicon were used by Hall-Stoodley et al. (2006) to study biofilm formation by *M. marinum*. *Mycobacterium chelonae, M. flavescens, M. fortuitum, M. gordonae, M. kansasii, and M. terrae* were found to form biofilms on organic substances such as plastics and rubber (Schulze-Röbbecke et al. 1992). Polyvinyl chloride (PVC) pipes were also used by Vess et al. (1993) to study biofilm growth of a range of bacteria including *M. chelonae*. Dailloux et al. (2003), Lehtola et al. (2007) and Torvinen et al. (2007) also demonstrated the attachment and accumulation of mycobacteria on HDPE, PVC and stainless steel surfaces. None of the research works reported here investigated biofilm formation and growth by Map which is an implicated agent of Crohn’s disease of humans, and the main cause of Johne’s disease of ruminants, hence an object for investigation in this report. Whilst biofilm could serve as a mode of transfer of Map in the water environment, also within and between both human and animal populations, this work was designed to firstly determine biofilm formation and growth of Map on PVC which is minimally employed in potable water production. This was however, preceded by an adhesion assay using aluminium and stainless steel which are also relevant materials in water treatment operations and storage devices. Secondly, the effect of schmutzdecke composites on biofilm formation by Map was also investigated.

## 2. MATERIALS AND METHODS

### 2.1 Preparation of aqueous extract of schmutzdecke and Map cultures

Aqueous extract of schmutzdecke [(AES; filtered (FAES) and unfiltered (UAES)], and distilled water (DW) were prepared as follows: The UAES was prepared by mixing 20 g schmutzdecke in 480 ml distilled water using Stomacher (Stomacher 400, Seward, England) for 4 min and allowing particles to settle (10 min) followed by centrifugation of the supernatant at 489 x g for 10 min to remove larger particles. The FAES was prepared by filtering a portion of the UAES through a 150 ml, 50 mm, 0.45 μm pore size filter unit (MF75™ Series, Nalgene, Fisher Scientific U.K. Ltd, Bishop Meadow Road, Loughborough, Leicestershire, LE11 0RG, UK). Both the filtered and unfiltered AES portions were irradiated at a dose of 25 kGy and were stored at 4°C until required. The already prepared DW (UKAS protocol) was also autoclaved at 121°C for 15 min and stored at 4°C until required. Three Map cultures containing 10^6^CFU ml^−1^ were prepared from working cultures of Map (strain ATCC 43015) and two water isolates (L 85 and L 87) with initial concentrations 10^8^ CFU ml^−1^ by centrifuging 1 ml of each working culture at 7558 x g for 20 min and reconstituting the pellets in 100 ml each of FAES, UAES and DW and determining the CFU ml^−1^ on Middlebrook 7H10 plates.

### 2.2 Adhesion by Map on aluminium and stainless steel coupons

Three 6-week old working cultures of Map (ATCC 43015, L 85 and L 87) were each reconstituted in PBS-T20 (1:100) to prevent clumping of cells. These were then incubated overnight in 30 mm Petri dishes in which were placed separately in a horizontal position sterile coupons made of aluminium and stainless steel (dimensions: 2 cm × 2 cm) for 48 hours at 37°C. Following the 2-day incubation period, the cells were washed 3 times with distilled water before placing the coupons on a heating block at 100°C for 2 min to heat-fix the cells. The coupons were then placed in Petri dishes (30 mm) containing 0.03% (w/v) acridine orange dye, and incubated at room temperature (approx. 21°C) for 20 min to allow penetration of the dye into the Map cell wall. The coupons were washed again with distilled water to remove remaining acridine orange dye before air drying and subsequent inspection using x 1000 magnification of a fluorescent microscope, and a photographic record was made.

### 2.3 Biofilm formation by Map

This procedure was followed for a period of 10 weeks. The media used were DW, FAES and UAES. Using 100 μl of 1 week old working cultures of Map (ATCC 43015, L 85 and L 87), selected wells of three 96-wells micro-titre plates made of polyvinyl chloride (PVC, flat bottom, Falcon 353912, Becton Dickinson and Company, Franklin Lakes, New Jersey, USA) were each inoculated with one isolate and then subjected to each of the media (DW, FAES and UAES) in separate wells to a final volume of 200 μl. The plates were sealed with parafilm and incubated at 37°C for a period of 10 weeks. The volume of the cultures were maintained by replenishing them with 100 μl of the respective media at weekly intervals over the entire period of the experiment. This was to mimic a similar environment in water treatment operations where slow sand filter beds receive known volumes of raw untreated water periodically to maintain the build-up of schmutzdecke. At the end of the 10 weeks, remaining media was removed from each well, and 200 μl of 1% (w/v) crystal violet (CV) was used to stain the cells for 30 min followed by three times of washing with the respective media. The optical density (OD) of cell-bound CV eluted into the wells after adding 200 µl of 95% (v/v) ethanol to the wells for 1 hour, was measured by means of a Spectrophotometer (Sunrise Remote, Tecan, Austria, Gmbh, 5082, Grodig, Austria) using a wavelength of 595 nm. This experiment was repeated in triplicate runs and the average OD_595nm_ values were calculated and plotted against time of incubation. The three runs for each isolate were meaned and the mean value used for statistical comparison both between isolates and over time.

## 3. RESULTS

### 3.1 Adhesion by Map on aluminium and stainless steel coupons

Figure 1 represents the adhered cells of Map on aluminium whereas Fig. 2 shows Map cells adhered to stainless steel. Both coupons possessed different surface properties (rough for aluminium and smooth for stainless steel). However, each of the 2 coupon types was of same surface properties (obtained from same individual batches of manufacture).

**Figs. 1 and 2:**
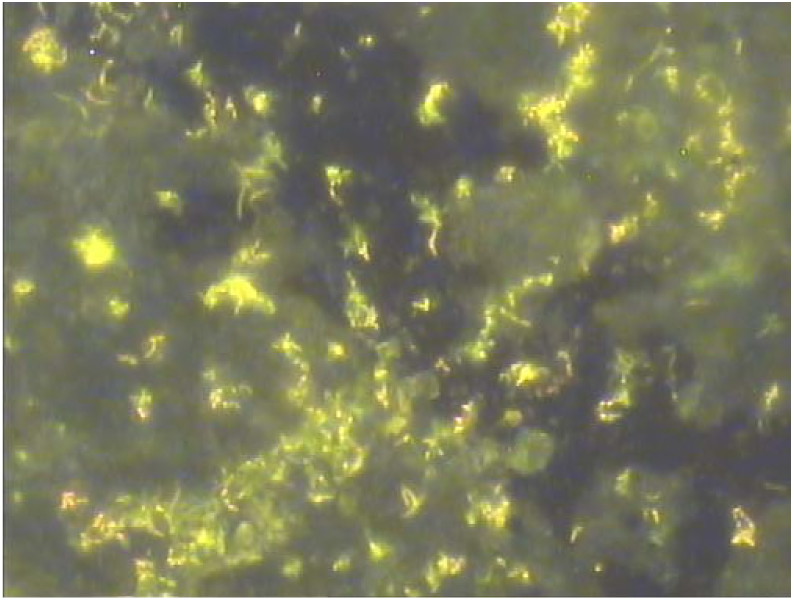
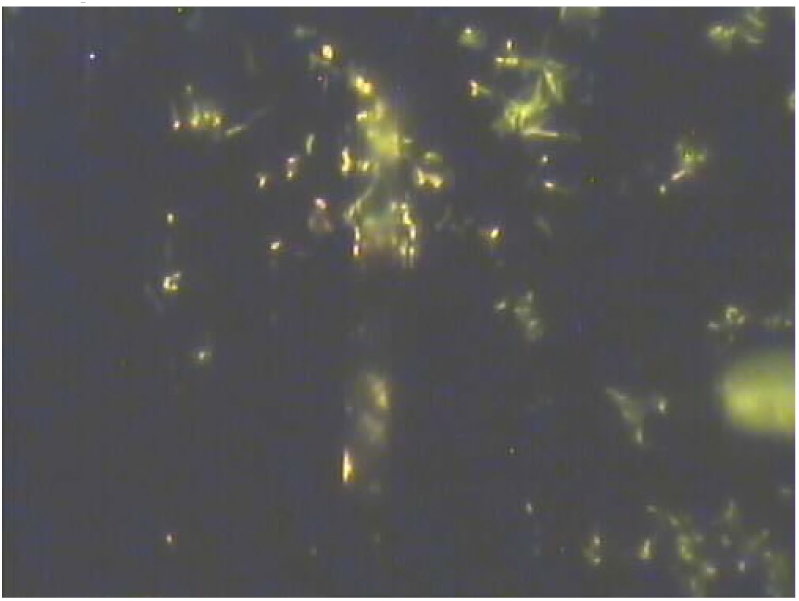
Adhesion of Map to aluminium (Fig. 1) and stainless steel (Fig. 2) coupons analysed and photographed by fluorescent microscopy after incubation in PBS-T20 for 2 days at 37°C.

**Fig. 3:**
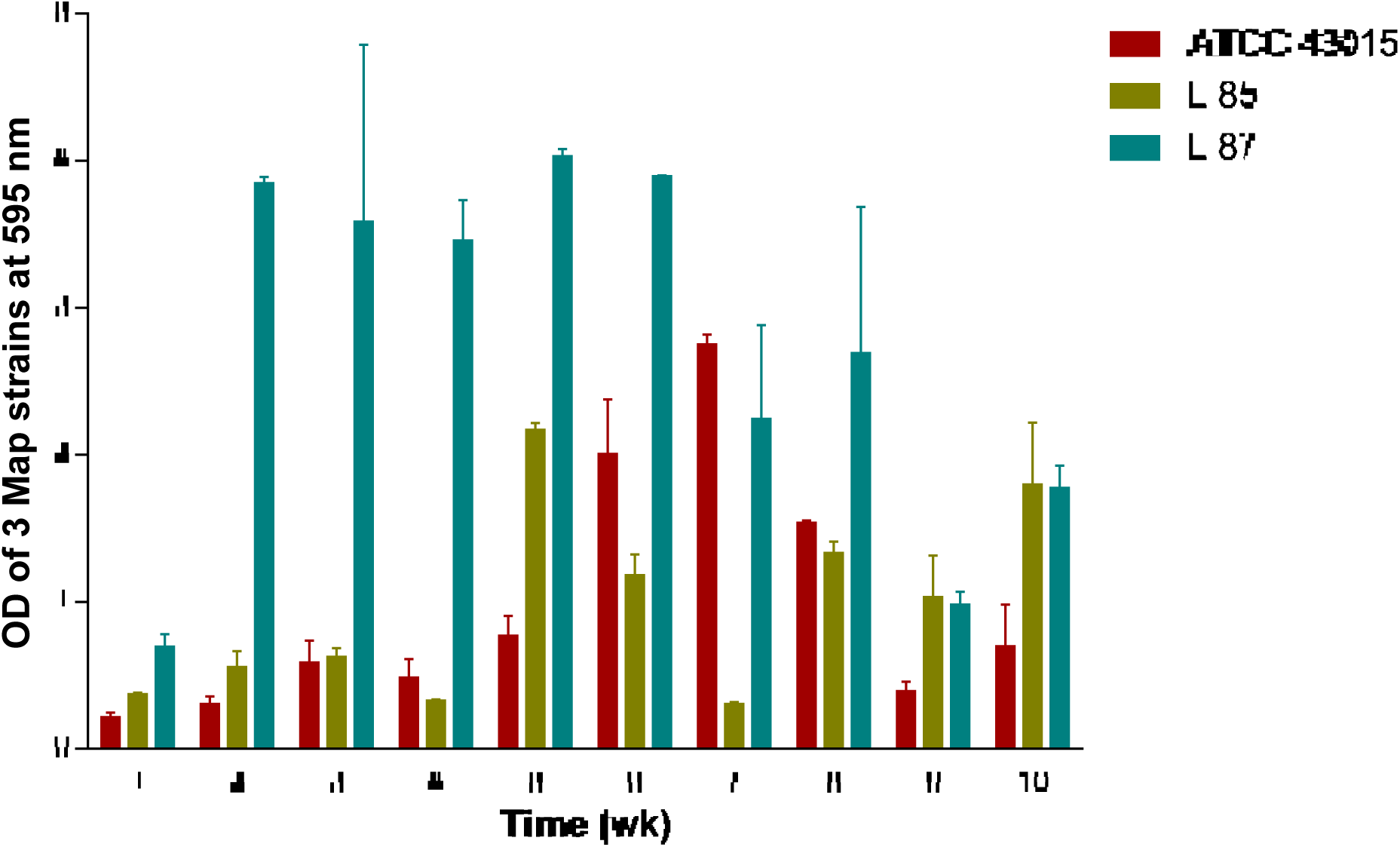
Biofilm formation and growth by 3 Map strains in DW (control)

**Fig. 4:**
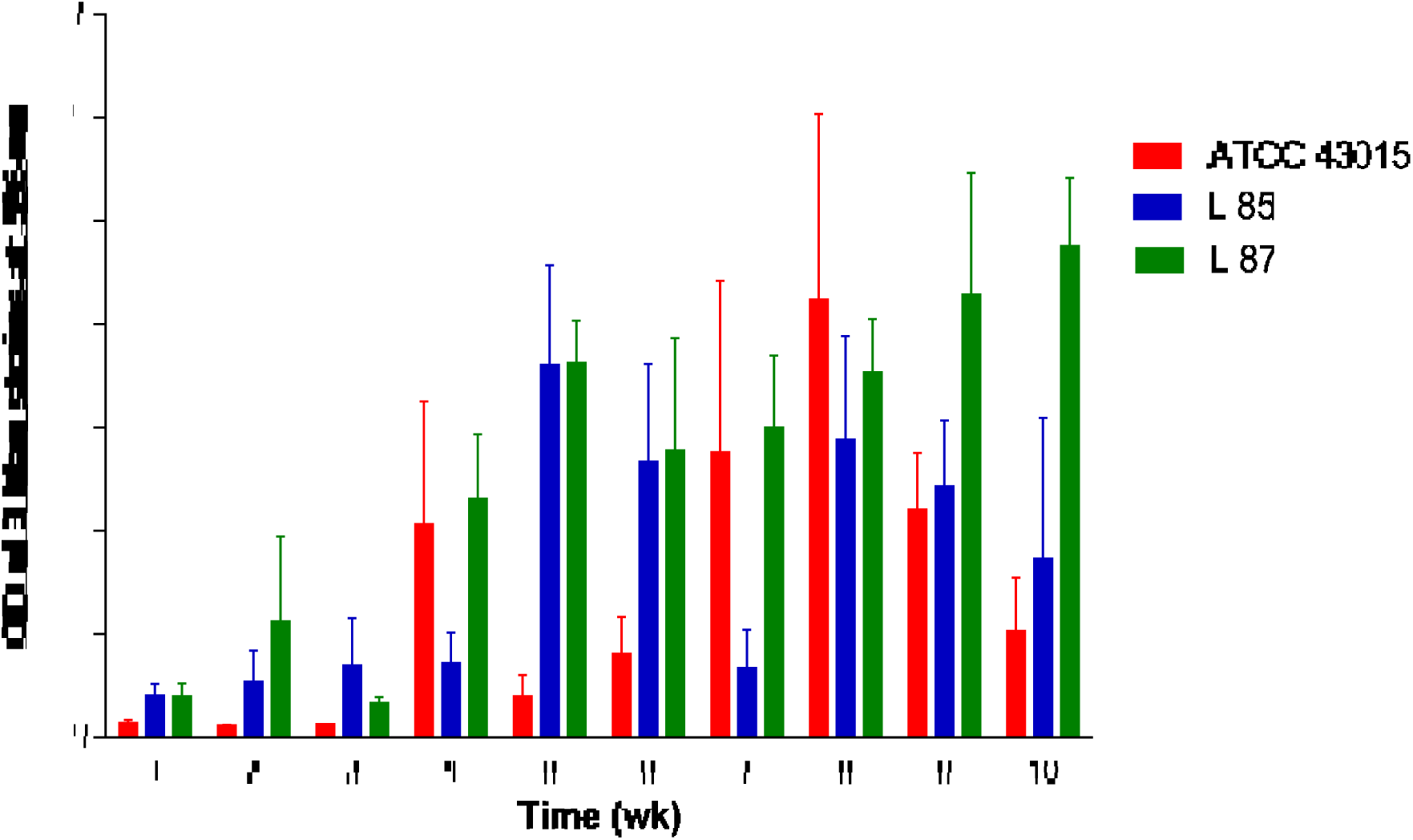
Biofilm formation and growth by 3 Map strains in FAES medium.

**Fig. 5:**
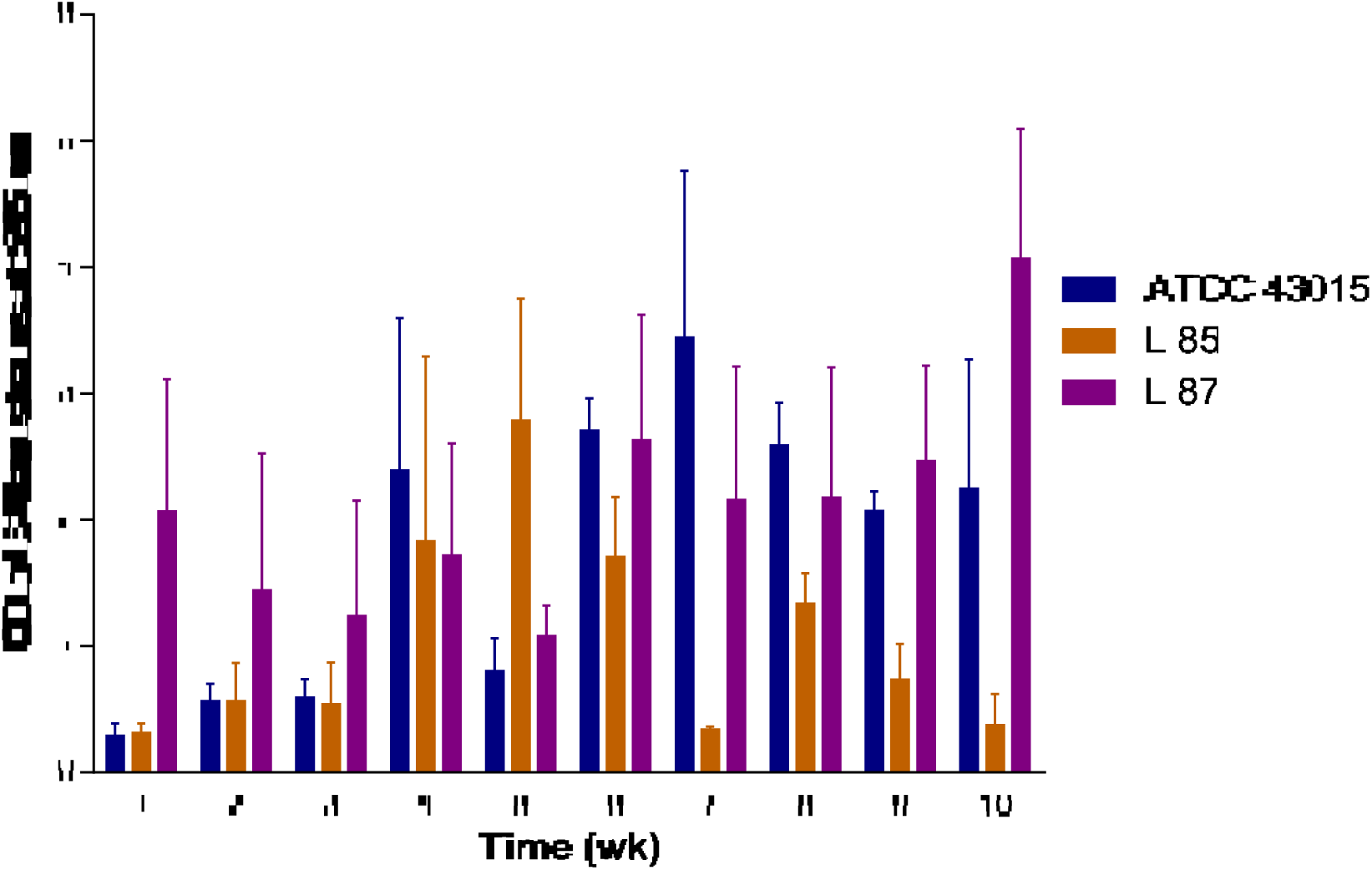
Biofilm formation and growth by 3 Map strains in UAES medium.

### 3.2 Biofilm formation by Map (ATCC 43015, L 85 and L 87) in DW (control)

An ANOVA showed a general significant difference (P < 0.05) in biofilm density amongst the 3 Map strains. The L 87 strain grew more cell mass compared to L 85 and ATCC 43015 strains. Thus L 87 had a better response to growth in DW than the other two strains in the first 8 weeks of incubation before dropping in cell density in the following weeks (9 and 10). The L 85 and ATCC 43015 pattern however, followed an irregular trend with a generally low cell density between week 1 to 8 compared with L 87.

### 3.3 Biofilm formation by Map (ATCC 43015, L 85 and L 87) in FAES

The ANOVA showed a significant difference (P < 0.05) amongst the 3 strains in their response to biofilm density in FAES medium over the 10-week period. The response here also showed an increasing response of biofilm density by L 87 than L 85 and ATCC 43015. Unlike in the UAES medium, L 85 improved in biofilm density in the FAES medium than the ATCC 43015 which responded less.

### 3.4 Biofilm formation by Map (ATCC 43015, L 85 and L 87) in UAES

An ANOVA showed a general significant difference (P < 0.05) in biofilm density amongst the 3 Map strains as influenced by UAES medium over the entire 10-week period. Even though the trend of response of the 3 strains to the growth medium was erratic, there was generally a greater increase in biofilm density by L 87 compared to ATCC 43015 and the least by L 85. Looking at the graph, strains ATCC 43015 and L 85 appear to have a sigmoid response; one peaking at week 7 and the other at week 5 with L 87 agreeing erratically. Thus in all the three media (DW, FAES and UAES), L 87 responded better followed by L 85 with ATCC 43015 responding less. It was noted that the water isolates (L 85 and L 87) grew denser biofilms in the aqueous extract of schmutzdecke than the Crohn’s disease isolate (ATCC 43015) depicting better performance from their originality as environmental isolates compared with ATCC 43015 as a human isolate.

## 4. DISCUSSION

### 4.1 Adhesion of Map to aluminium and stainless steel coupons

Although there is only minimum use of aluminium or stainless steel in water pipes and water treatment works they were chosen as substrata for laboratory-based biofilm studies. The ATCC 43015 strain of Map was also chosen for this work as the reference strain partly due to the fact that it was obtained from a human source and has also been cultivated in the laboratory over a period, hence considered as having a wider adaptation to different environments. There is no doubt that mycobacteria form biofilm; examples are *M. kansasii* (Schulze-Röbbecke, 1993), *M. flavescens* (Schulze-Röbbecke and Fischeder, 1989) and *M. ulcerans* (Marsollier et al. 2005). However, to the author’s knowledge there is no published information on Map forming a biofilm under environmental conditions. *Mycobacterium kansasii* and *M. marinum* are slow growing and pathogenic like Map while *M. flavescens* is fast growing and non-pathogenic. The fact that particularly the slow growing *Mycobacterium* spp. form biofilm provides evidence that at least Map was likely to associate, if not grow in biofilm. Also there is circumstantial evidence that a close relative of Map i.e. *M. avium* ssp. *avium* can form biofilm in that it has been reported to persist in actively flowing streams (Primm et al. 2004) presumably as a result of forming biofilm on the rocks present. In the absence of direct evidence of the ability of Map to form biofilm, an experiment was undertaken to determine if Map could adhere to aluminium and stainless steel coupons which would represent the initial stages of biofilm formation and production. It is clear that the isolates adhered to both surfaces after 2 days of incubation. In particular, there was a greater density of cells on the aluminium surface. The aluminium coupon had a much rougher surface and it is unclear to what extent this afforded protection against the shear force of the wash water. Also the rough surface may have allowed more of the cells outer wall to come into contact with the surface and may have allowed more cells to be attached to the aluminium coupon than the stainless steel coupon.

### 4.2 Biofilm formation by Map

Following research reports that mycobacteria could form biofilm, and success of the adhesion test with the Map isolates, it seemed prudent to determine biofilm formation by Map, and the extent to which this was a common trait amongst its isolates as these were not reported in literature. Whilst 37°C provided optimum growth temperature for Map, the question as to whether the three isolates would behave or grow optimally when environmental components of water treatment works were also present needed to be answered. The environmental components chosen were DW as control, FAES and UAES. The two Map isolates (L 85 and L 87) showed better growth in both the FAES and UAES over the ATCC 43015 strain, perhaps to be expected since they are environmental isolates and may have little or no nutrients in the environment compared to the biofilm medium. This is consistent with the observation that with no added nutrients, *M. avium* grew in water medium (George et al. 1980). However, the FAES and UAES media suspensions are rich in both humic and fulvic acids which partly serve as metabolic nutrients for environmental *M. avium* complex in the coastal areas (Falkinham 1996; Kirschner et al. 1999). Since the L 85 and L 87 strains were obtained from the water environment, they might have derived and better assimilated these metabolic nutrients than the ATCC 43015 strain, agreeing to a possible variation in biofilm formation by Map isolates (Johansen et al. 2009), with the ATCC 43015 peaking growth around week 7 in all the three media. It can be seen that all the three strains of Map showed optimum growth after 5 weeks, however, the decline in growth following this duration of time, may be attributed to exhaustion of nutrients and production of metabolites that were inhibitory to growth. However, L 87 generally had more biofilm growth in DW between weeks 1-6 and also in both FAES and UAES weeks 9-10 compared to the L 85 strain which also grew a denser biofilm than the ATCC 43015 strain. It is worth commenting that there was generally no difference in growth response between FAES and UAES. Aside from the role of the 3 media in biofilm growth of the three Map strains, the make-up of the PVC plate employed for this work may also have contributed to the biofilm formation by the 3 strains, by providing a suitable substratum (Norton et al. 2004) since more substantive mycobacterial biofilm have been shown to occur on plastics and rubber compared to other substrata (Schulze-Röbbecke et al. 1992). The findings from this research work thus demonstrate the degree by which survival and proliferation of Map is influenced by the composition of its environment.

## 5. CONCLUSIONS

The formation of biofilm by Map is influenced by the properties of its environment. The water environment serves as a possible hub of Map’s persistence in a biofilm which is compounded by variation of its nutrient composition, substratum and prevailing conditions. Irrespective of the originality and source of isolation, and also traits and differences amongst strains, Map adapts to biofilm formation and growth subject to prevailing conditions. Incorporation of smooth and/or biofilm-unfriendly surfaces in water treatment operations and storage devices can reduce or curtail the formation of biofilm by Map and its spread in the water environment.

## ACKNOWLEDGEMENTS

The authors wish to express their gratitude to Queen’s University Belfast, for funding this research, and Agri-Food and Biosciences Institute for allowing access to their laboratories.

